# Structural basis for the ARF GAP activity and specificity of the C9orf72 complex

**DOI:** 10.1101/2021.04.13.439632

**Authors:** Ming-Yuan Su, Simon A. Fromm, Jonathan Remis, Daniel B. Toso, James H. Hurley

## Abstract

Mutation of *C9ORF72* is the most common genetic cause of amyotrophic lateral sclerosis (ALS) and frontal temporal degeneration (FTD), which is attributed to both a gain and loss of function. C9orf72 forms a complex with SMCR8 and WDR41, which was reported to have GTPase activating protein activity toward ARF proteins, RAB8A, and RAB11A. We determined the cryo-EM structure of ARF1-GDP-BeF_3_^-^ bound to C9orf72:SMCR8:WDR41. The SMCR8^longin^ and C9orf72^longin^ domains form the binding pocket for ARF1. One face of the C9orf72^longin^ domain holds ARF1 in place, while the SMCR8^longin^ positions the catalytic finger Arg147 in the ARF1 active site. Mutations in interfacial residues of ARF1 and C9orf72 reduced or eliminated GAP activity. RAB8A GAP required ∼10-fold higher concentrations of the C9orf72 complex than for ARF1. These data support a specific function for the C9orf72 complex as an ARF GAP.

## Introduction

Expansion of GGGGCC repeats in the first intron of *C9ORF72* is the most prevalent genetic cause of two clinically overlapping neurodegenerative diseases, amyotrophic lateral sclerosis (ALS) and frontal temporal degeneration (FTD) ^1,2^. There are two mutually non-exclusive mechanisms for *C9ORF72* expansion-induced pathogenesis in ALS/FTD, gain of function from the presence of toxic RNA and dipeptide repeats ^3,4^, and loss of function due to reduced C9orf72 protein expression ^5-7^. The reduction or absence of C9orf72 has been proposed to alter vesicle trafficking ^8- 10^, autophagy ^10-13^, lysosomal homeostasis ^14-16^, actin dynamics ^17^ and inflammation ^18^. However, the direct biochemical function(s) of C9orf72 have been uncertain until recently.

C9orf72 exists in cells as a stable heterotrimer with Smith–Magenis chromosome regions 8 (SMCR8) and WD repeat-containing protein 41 (WDR41)^9,10,12-15^. Both C9orf72 and its cellular binding partner SMCR8 are longin and DENN domain (differentially expressed in normal and neoplastic cells) containing proteins ^19^. Two cryo-electron microscopy structural studies of the C9orf72:SMCR8:WDR41 complex found that SMCR8 is sandwiched between C9orf72 and WDR41 ^20,21^. Structural similarity was observed between C9orf72:SMCR8 and another double longin-DENN complex, the folliculin and folliculin-interacting protein 2 (FLCN-FNIP2) complex ^22,23^. FLCN-FNIP2 is a GTPase activating protein (GAP) complex for RagC in the mTORC1 signaling pathway, with the putative catalytic Arg residue residing in the FLCN subunit. This suggested that C9orf72:SMCR8 might also be a GAP for some small GTPase ^20,21^. The C9orf72 complex was reported to have GAP activity for both ARF family proteins ^21^, and the RAB proteins RAB8A and RAB11A ^20^. The conserved Arg147 of SMCR8 was proposed to serve as the catalytic Arg finger typical of most GAP proteins. Mutation of SMCR8 Arg147 eliminated GAP activity, consistent with the Arg finger hypothesis.

The discovery that the C9orf72 complex has GAP activity against subsets of both ARF and RAB proteins was exciting in that both of these small GTPase families participate extensively in membrane traffic ^24^, as does C9orf72 ^5,8,25^. ARF6 and RAB11 vesicles^26,27^ have been implicated in retrograde vesicle transport in axons, such that excessive ARF6^GTP^ and RAB11^GTP^ have been proposed to inhibit anterograde transport and axonal regeneration^28^. Nevertheless, the reported dual specificity of the C9orf72 complex is unusual. ARF and RAB proteins belong to distinct small GTPase families. Yet C9orf72 is inactive as a GAP against the close ARF relatives ARL8A and B, and also inactive against RAB1A and 7A ^21^. Thus, the C9orf72 complex is not simply a promiscuous GAP. In order to clarify the specificity of the C9orf72 complex GAP activity, we sought to determine structures of the complexes with representative ARF in GAP-competent states. We selected ARF1 as a structurally well-characterized model for the ARF family, having previously found that GAP activity against ARF1, 5, and 6 was indistinguishable ^21^. We found that the GAP reaction went to completion at an order of magnitude lower C9orf72 complex concentration for ARF1 as compared to RAB8A (> 10-fold). Consistent with a preference for ARF proteins, it was possible to determine the structure of the ARF1:C9orf72:SMCR8:WDR41 complex in the presence of the GTP hydrolysis ground state mimetic BeF_3_^-^. These data validate that the ARFs are specific substrates of C9orf72 complex GAP activity.

## Results

We first characterized the concentration of C9orf72:SMCR8:WDR41 needed to produce complete hydrolysis of GTP by ARF1. An ARF1 construct with the N-terminal myristoylated amphipathic helix removed was used in all experiments, as this region was previously found to be unnecessary for its competence as a GAP substrate ^21^. The concentration of ARF1 was fixed at 30 µM, and then treated with 0, 0.3, 1.5 and 3 µM C9orf72 complex for 15 min at 37°C, and nucleotide products were monitored by high pressure liquid chromatogram (HPLC) analysis. The 0.3 µM C9orf72:SMCR8:WDR41 sample, equivalent to a 1/100 molar ratio relative to ARF1, was sufficient to completely hydrolyze all of the GTP bound to ARF1 (Extended Data Fig 1a).

**Fig.1:**
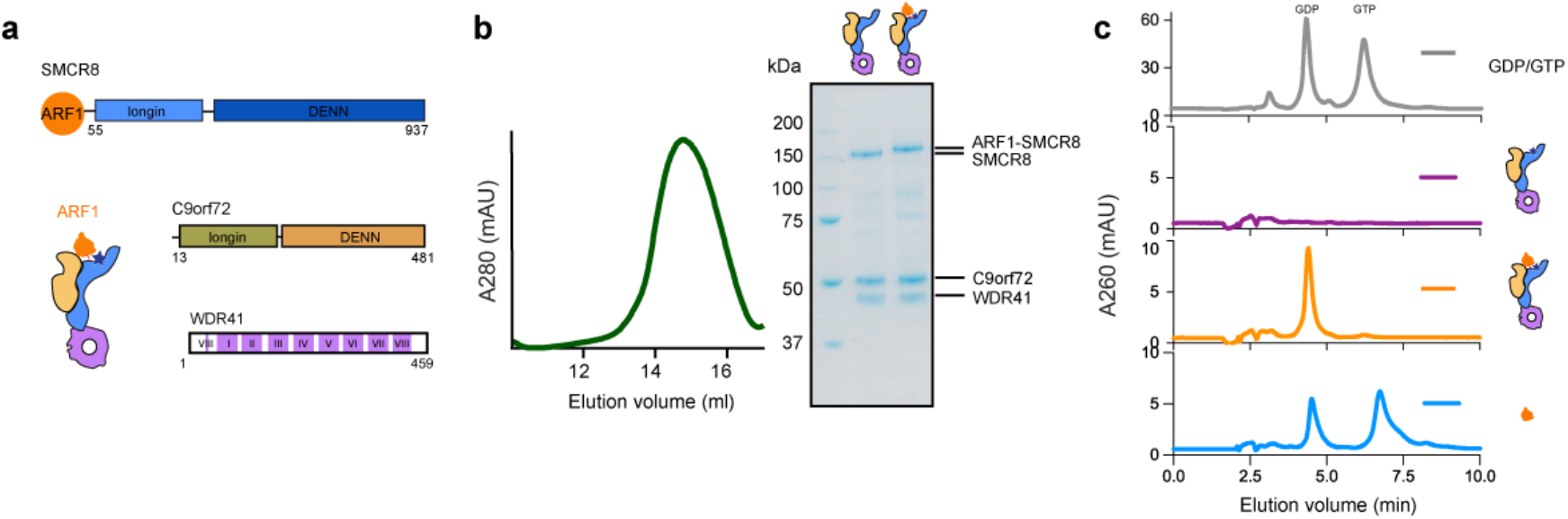
Construct design of C9orf72:ARF1-SMCR8:WDR41 complex. a, Diagram of the complex design. ARF1 is fused to the N-terminus of SMCR8 subunit. The blue star represents the position of the catalytic Arg residue of SMCR8. b, Left, the Superose 6 size exclusion profile of the reconstituted ARF1-SMCR8:C9orf72:WDR41 complex. Right, the SDS-PAGE analysis of the peak fractions for ARF1-SMCR8:C9orf72:WDR41 complex compared with wild type complex. c, The HPLC analysis of the purified C9orf72:SMCR8:WDR41, C9orf72:ARF1-SMCR8:WDR41 complex and ARF1. The elution profile for GDP/GTP is shown on the top panel.

To probe the structural basis of the catalytic mechanism, we attempted to assemble the GAP-ARF1 complex by mixing the GTP-locked mutant ARF1^Q71L^ with the C9orf72:SMCR8:WDR41 protein complex. A stable complex was observed by gel filtration chromatography (Extended data Fig. 2a,b), but not by cryo-EM (Extended data Fig. 2c). We inferred that the complex dissociated during vitrification. Following a precedent in which it was found necessary to use a fusion construct to trap the structure of the ASAP3:ARF6 GAP complex ^29^, we fused the C terminus of ARF1 to the flexible N-terminus of SMCR8, and coexpressed the fusion with C9orf72 and WDR41 in HEK293 GnTi cells as a chimeric C9orf72:ARF1-SMCR8:WDR41 complex (Fig 1a). The N-terminal 50 residues of SMCR8 are disordered ^21^, effectively serving as a flexible linker between ARF1 and SMCR8 of up to 150 Å in length, which is similar to the longest axis of the C9orf72:SMCR8:WDR41 complex. The size exclusion profile demonstrated a similar yield and hydrodynamic behavior to the wild type complex (Fig 1b). HPLC analysis was used to assay the identity of the nucleotide bound in ARF1. Recombinant ARF1 protein as isolated from cells was 45-55 % GTP loaded, whereas the fusion version was only GDP-bound (Fig. 1c). This suggested to us that the fusion construct was catalytically active.

We generated a nucleotide hydrolysis ground state analog ^30^ by incubating the GDP-bound protein complex with BeF_3_^-^ overnight, and visualized it by cryo-EM. A preliminary reconstruction of C9orf72:ARF1-SMCR8:WDR41 was obtained from a dataset collected on a 200 kV Talos Arctica. This reconstruction showed extra density corresponding to ARF1 localized atop the two longin domains (Extended Data Fig. 3), and coordinates for the GTP-bound ARF1 model (PDB 1O3Y) ^31^ fit the density. However, some regions were poorly resolved, and additional data sets were collected on a 300 kV Titan Krios microscope (Extended data Fig. 4-6). We were able to obtain a 3.9 Å cryo-EM reconstruction from an untilted dataset combined with 30° and 40° stage tilted datasets. It was possible to visualize the locations of the bound nucleotide and larger side-chains in this reconstruction, but smaller side-chains were not resolvable, nor were conformational details of larger side-chains.

The reconstruction showed that ARF1 sits in a 25 Å-wide concave pocket generated by the SMCR8^longin^ and C9orf72^longin^ domains. The ARF1 switch I and switch II regions, which are conformationally sensitive to the nucleotide state, directly contact the parallel helices of the SMCR8^longin^-C9orf72^longin^ dimer (Fig. 2a,b). The bowl formed by the two longin domains can be compared to the bowl of a spoon, and the parallel N-terminal helices of SMCR8 to the spoon handle. Relative to C9orf72, SMCR8^longin^ has the more extensive interface with ARF1 (Fig. 2b). ARF1 is in its GTP bound conformation, with respect to all three of the regions whose conformation depends on the nucleotide state: the switch I, switch II, and interswitch regions (Fig. 2c).

**Fig.2:**
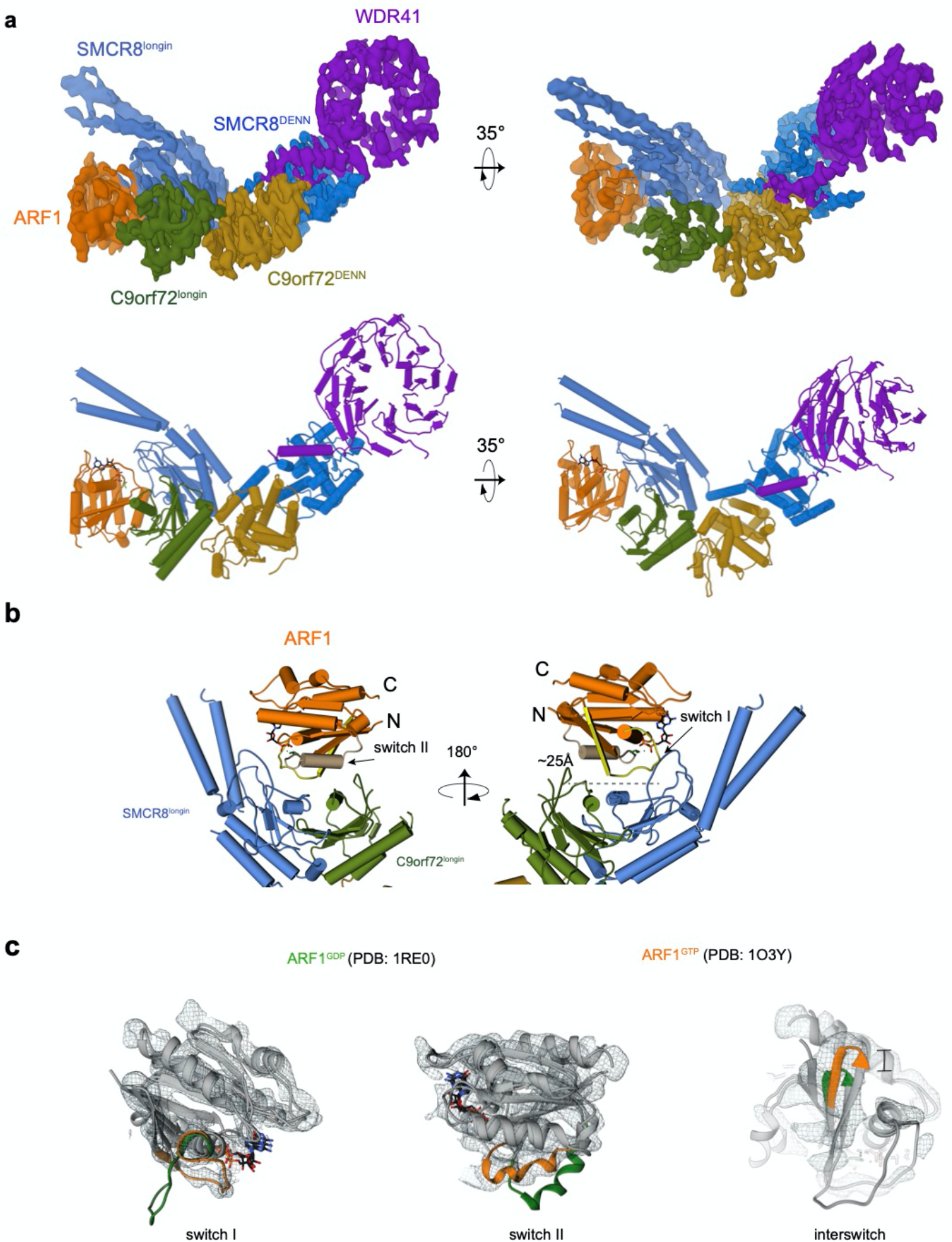
Cryo-EM structure of C9orf72:ARF1-SMCR8:WDR41 complex. a, Cryo-EM densities and the refined atomic model of the complex. The domain colored as followed: SMCR8^longin^, cornflower blue; SMCR8^DENN^, dodger blue; C9orf72^longin^, olive; C9orf72^DENN^, goldenrod; WDR41, medium purple; ARF1: orange. b, ARF1 sits on the top of the two longin domains. The switch I and II of ARF1 are colored in yellow and tan, respectively. c, Structural comparison of ARF1 in GTP versus GDP bound states (from the left to right panels, switch I and switch II and interswitch). ARF1 in GTP state fits into the density.

SMCR8 Arg147 points directly towards the BeF_3_^-^, as predicted by the arginine finger model (Fig. 3a, b, left). The geometry is similar to that seen in the only other structure of a catalytically competent ARF GAP complex, that of ASAP3 and ARF6-GDP-AlF_x_ (PDB: 3LVQ) ^29^ (Fig. 3a, b, middle). In the ASAP3:ARF6 structure, only one face of ARF6 contacts the GAP. In comparison, the C9orf72-SMCR8 longin spoon bowl cups ARF1 more extensively. ARF1 switch II sits on the parallel helices of the two longin domains and the loop of SMCR8^longin^, while switch I binds only to SMCR8^longin^. The more extensive cupping across the Switch I/II regions is seen in other GAP complexes, however, such as the TBC1D20:RAB1B-GDP-BeF_3_^-^ complex (PDB: 4HLQ) ^32^ (Fig. 3a, b, right).

**Fig.3:**
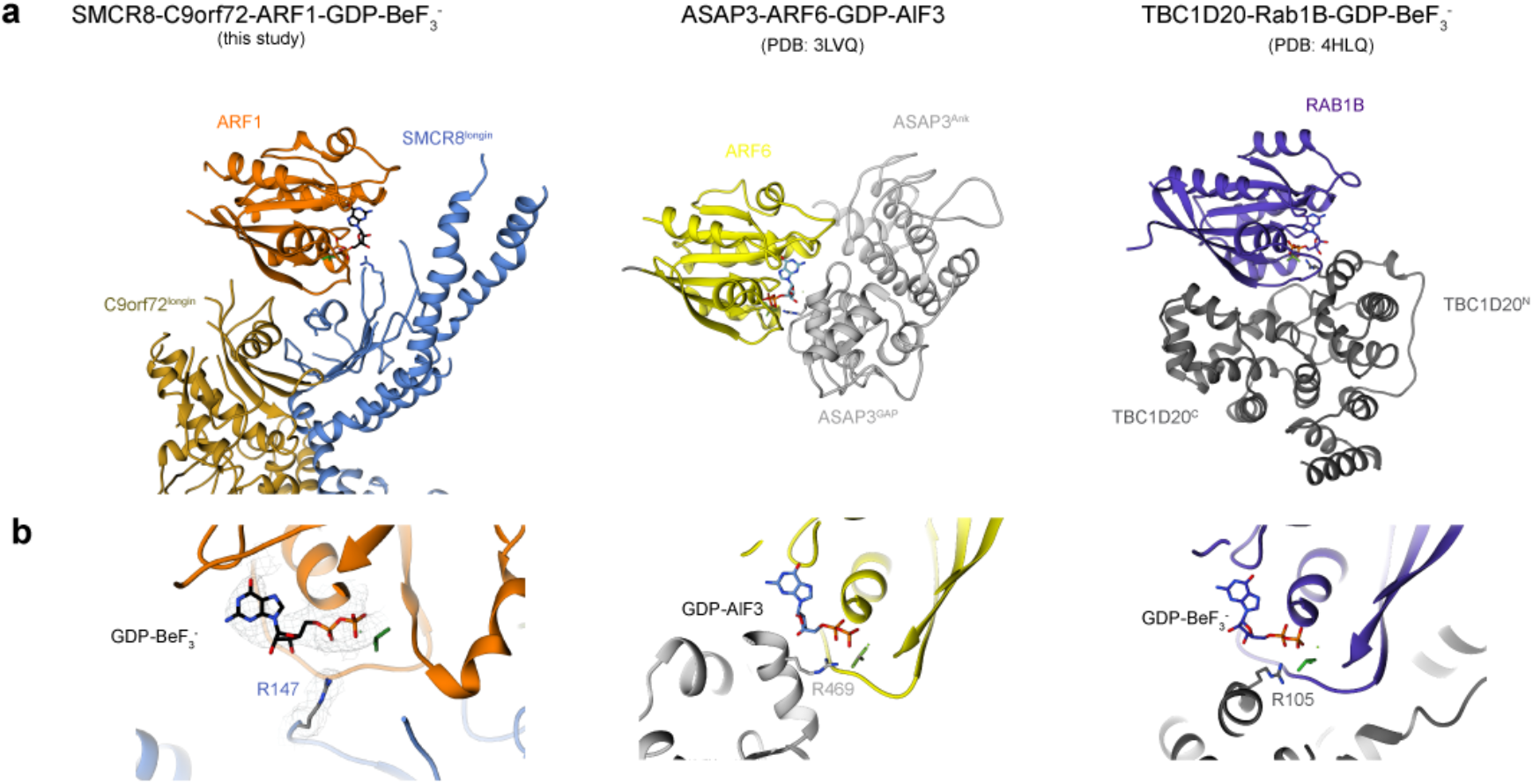
Structure comparison between C9orf72:ARF1-SMCR8:WDR41 with other GAP:small GTPases. a, Structure comparison between C9orf72:ARF1-SMCR8:WDR41 (left) with ASAP3:ARF6 (middle) and TBC1D20:RAB1B (right). The small GTPases are shown in the same orientations. b, Close-up views of the nucleotide binding sites with catalytic Arginine finger residues pointing to the active sites.

ARF1 residues at the C9orf72 and SMCR8 interface were mutated and tested for GAP activity as monitored by HPLC (Fig. 4a, b). Mutation of residues (I49H, G50R) located at the switch I region largely reduced the activity compared to the ones at the switch II region (K73W), which is in consistent with the more extensive interface of SMCR8. We mutated the C9orf72 interface residues Trp33 and Asp34 in the double mutant W33G/D34K (Fig.4c). C9orf72^W33G/D34K^:SMCR8:WDR41 complex was found to almost completely abolish GAP activity towards ARF1. We confirmed that C9orf72:SMCR8 was active towards RAB8A, albeit at much higher concentrations of the GAP complex than for ARF1. GAP activity with respect to RAB8A requires a 1/10 molar ratio of C9orf72:SMCR8:WDR41 to hydrolyze the RAB8A proteins, at least ∼10-fold more than for ARF1 (Extended Data Fig. 1b). The I71R mutation of Ile71 on the C9orf72^longin^ helix decreases the activity towards ARF1, whilst essentially abolishing activity towards RAB8A (Fig. 4).

**Fig.4:**
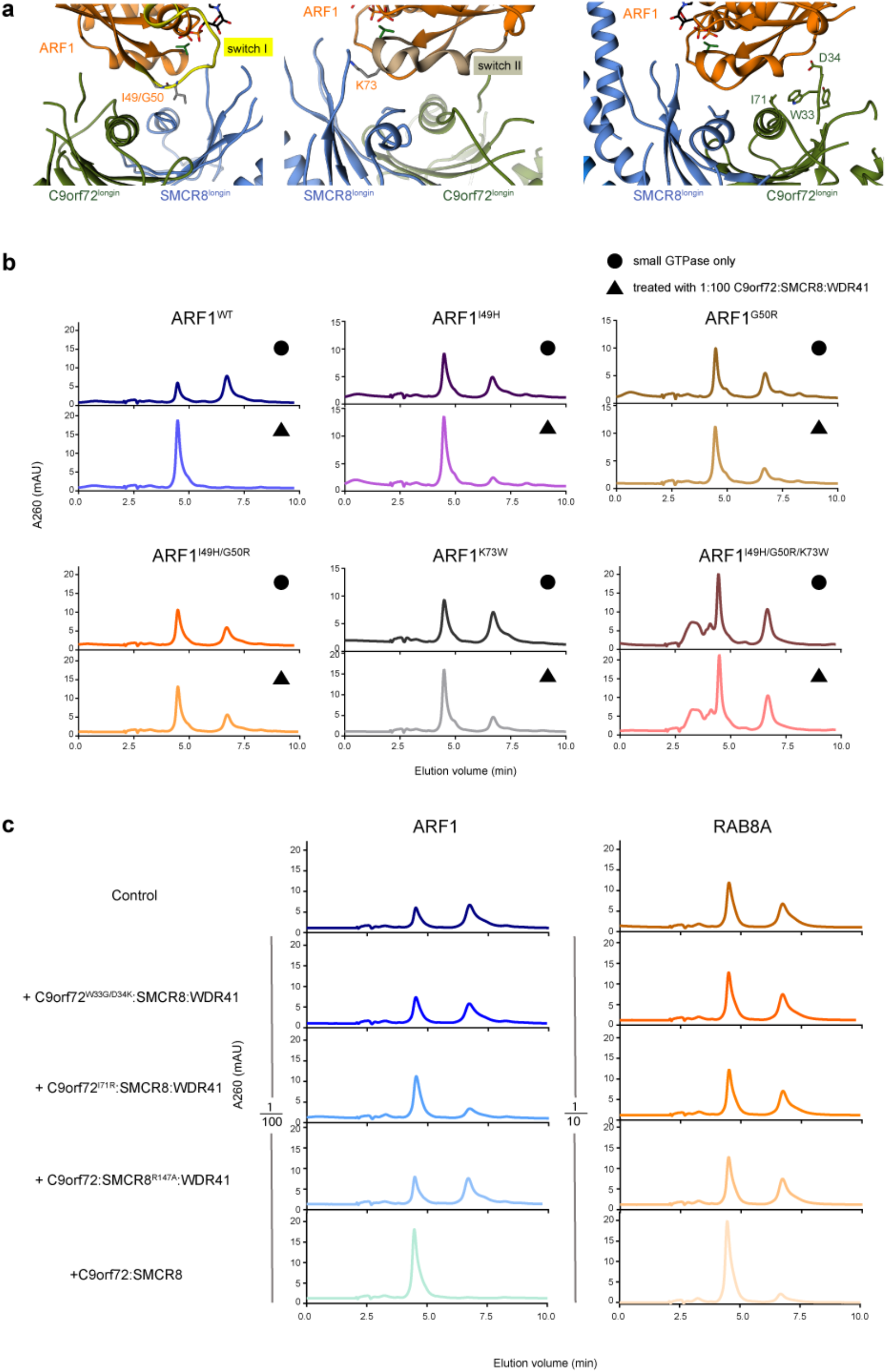
Interfaces between ARF1 and C9orf72/SMCR8 and HPLC result of ARF1^WT/mutants^ with C9orf72^WT/mutant^:SMCR8:WDR41. a, Close up views of the interfaces between switch I/switch II regions of ARF1 with C9orf72:SMCR8. b, HPLC analysis of ARF1^WT/mutants^ incubated with C9orf72:SMCR8:WDR41. The top panels are the control reactions for each ARF1 proteins, either wild-type or mutants. The bottom ones are incubated with 1/100 molar ratio of C9orf72:SMCR8:WDR41 c, ARF1 or RAB8A incubated with 1/100 or 1/10 molar ratio of C9orf72^WT/mutants^:SMCR8:WDR41, respectively. The experiments in b, c are carried out at least three times and one representative plot is shown.

## Discussion

C9orf72-SMCR8 is one of a class of dimeric longin-DENN domain complexes that also include the FLCN-FNIP complex. Catalytic Arg residues of GATOR1 subunit Nprl2 ^33^, FLCN ^22,23^, and SMCR8 ^20,21^ have been proposed to mediate GAP activity by an Arg finger mechanism ^34^. We have now trapped and visualized the ARF1 GAP complex of C9orf72:SMCR8, confirming that the proposed Arg 147 residue of SMCR8 serves as a catalytic Arg finger in the expected geometry. Thus far it has not been possible to trap GATOR1 or FLCN-FNIP in catalytically competent GAP complexes with their substrates RagA and RagC, owing to the transient nature of the complexes and competition with more stable inhibitory states. The structure provided here will provide a basis for more precise modeling of the GATOR1 and FLCN GAP complexes, and suggests tactics that may make it possible to trap these structures as well.

The great majority of the >30 known ARF GAP proteins contain a conserved ARF GAP domain, first described for ArfGAP1 ^35^. Just a handful of ARF GAPs function without a canonical ARF GAP domain, including SEC23, the GAP for the ARF-related SAR1 GTPase, and the ELMO domain proteins ELMOD1-3 ^35^. The data presented above define the C9orf72:SMCR8 longin-DENN complex as a new addition to the small set of protein folds possessing ARF GAP activity. We previously found that C9orf72:SMCR8 was active against ARF1, ARF5, and ARF6, although it had no activity against ARL8A and B ^21^. The epitope immediately surrounding the GTP-binding site of ARFs 1-6 is highly conserved, and GAPs and GEFs are therefore typically active across the family ^36^. ARF6 regulates axon extension ^37^, and dominant-negative ARF6 expression rescues the axonal actin dynamics phenotype in *C9ORF72* ALS patient cells ^17^. While ARF6 is the best candidate for a C9orf72 GAP substrate relevant to neurodegeneration, the structural features of the ARF1 complex presented here are expected to be essentially identical between the two.

Our data strongly support that C9orf72:SMCR8 is a GAP for ARF GTPases. Given the report that C9orf72:SMCR8 is also a GAP for RAB8A and RAB11A ^20^, the question as to what are the physiological substrates of C9orf72:SMCR8 has become important in C9ORF72, ALS, and FTD research ^38,39^. The data presented here affirm C9orf72:SMCR8 as a *bona fide* ARF GAP by presenting a well-ordered cryo-EM structure which makes verifiable predictions of the activity of point mutants of both ARF1 and C9orf72. The generally much lower activity measured for RAB8A leaves the functional significance of the RAB GAP activity uncertain.

Another difference between the structures presented here and previously ^21^ is that our complex, expressed in HEK 293 cells was monomeric, while material expressed in insect Sf9 cells was dimeric ^20^. We noticed a small population of C9orf72:SMCR8:WDR41 dimeric complex in our cryo-EM sample, and the dimeric complex was more populated when we cross-linked it at high concentrations of protein complex. This suggests that that there is monomer-dimer equilibrium in the solution. Our observation of a catalytically competent monomeric complex does show that the monomer is the minimal entity capable of GAP activity.

The C9orf72 complex is reversibly localized to lysosomes in amino acid starvation through its WDR41 subunit ^40^. WDR41 in turn binds to the lysosomal cationic amino acid transporter PQLC2 ^41^. The PQLC2 binding site on WDR41 was recently identified ^42^, and is near the distal tip of the overall complex with respect to the ARF binding site. It is not apparent, based on existing structural data, how C9orf72:SMCR8:WDR41 could be localized to the lysosomal membrane to carry out GAP activity in *cis* on a small GTPase localized to the same membrane. There are few reports of ARFs localized to lysosomes, and thus is would not be surprising that the complex is not designed to act in *cis* while lysosomally localized. This leaves two models for lysosomal regulation of ARF GAP activity. In the sequestration model, C9orf72 is active as a GAP in amino acid replete cells, but inactivated by lysosomal sequestration in starved cells. Alternatively, in the *trans*-activation model, the ARF GAP active site could project away from the lysosome to act on a pool of ARF localized to nearby membranes or cytoskeletal elements under starvation conditions.

The working hypothesis suggested by the data above, is that C9orf72 haploinsufficiency in ALS and FTD results in an excess of ARF^GTP^ in one or more localized pools within the cell. These might be in the vicinity of lysosomes and/or within axons. The locations and natures of these pools are major open questions whose answers call for further study. Given the numerous roles of ARF proteins in membrane traffic ^35^, it would not be surprising if excess ARF^GTP^ led to multiple defects in signaling, membrane traffic, and autophagy, which are correlates of *C9ORF72* haploinsufficiency in disease ^6,7^. This idea does suggest a potential therapeutic approach for ALS and FTD. In principle, ARF^GTP^ levels could be rebalanced by inhibiting the ARF GEFs responsible for the GTP loading of the relevant local pools of ARF^GTP^, if the appropriate ARF GEF(s) were identified. This family of ARF activators comprises at least 15 members ^35^, which are the targets of the relatively unspecific natural product brefeldin A. A number of highly specific ARF GEF inhibitors have been identified, and strategies for their screening are well worked out ^43^, lending support to the plausibility of such an approach.

## Acknowledgments

We thank Roberto Zoncu and Aaron Joiner for discussions, and Richard Hooy for the workstation support. This work was supported by Department of Defense Peer Reviewed Medical Research Program Discovery Award W81XWH2010086 (J.H.H.), a postdoctoral fellowship from the Association for Frontotemporal Degeneration (M.-Y.S.), and an EMBO Long-Term postdoctoral fellowship (S.A.F.).

## Author contributions

M.-Y.S. and J.H.H. conceived the overall project, designed the study and wrote the original draft. M.-Y.S. designed and carried out all the experiments. S.A.F. advised on experimentation and validated the model. J.R. and D.B.T. assisted in cryo-EM data collection. All authors contributed to reviewing and editing the manuscript.

## Data availability

EM density map has been deposited in the EMDB with accession number EMD-23827. Atomic coordinates have been deposited in the PDB with accession number 7MGE.

## Material and methods

### Protein expression and purification

The gene encoding ARF1 (residues E17-K181) was amplified and inserted into a pCAG vector encoding an N-terminal STREP-STREP-FLAG tagged SMCR8 using KpnI restriction sites, resulting in a STREP-STREP-FLAG-ARF1-SMCR8 construct. The pCAG vector encoding an N-terminal glutathione *S*-transferase (GST) followed by a TEV cleavage site was used for expression of C9orf72. WDR41 was cloned into a pCAG vector without a tag. The complex was expressed in HEK293-GnTi cells as described previously ^21^. The gene encoding RAB8A (residues M1-S181) was subcloned into pCAG resulting in a STREP-STREP-FLAG-RAB8A. All the ARF1 mutants were created by site-directed mutagenesis using KAPA HiFi HotStart ReadyMix. The C9orf72 mutants were generated by two step PCR and ligated into the pCAG-GST vector.

HEK293-GnTi cells adapted for suspension were grown in Freestyle media supplemented with 1% FBS and 1% antibiotic-antimycotic at 37 °C, 80 % humidity, 5% CO_2_, and shaking at 140 rpm. Once the cultures reached 1.5–2 million cells mL^−1^ in the desired volume, they were transfected as follows. For a 1 L transfection, 3 mL PEI (1 mg ml^−1^, pH 7.4, Polysciences) was added to 50 mL hybridoma media (Invitrogen) and 1 mg of total DNA (isolated from transformed *E. coli* XL10-gold) in another 50 mL hybridoma media. Each 1 mg of transfection DNA contained an equal mass ratio of the three C9orf72 complex subunit expression plasmids. PEI was added to the DNA, mixed and incubated for 15 min at room temperature. 100 mL of the transfection mix was then added to each 1 L culture. Cells were harvested after 3 days. Cells were lysed by gentle rocking in lysis buffer containing 50 mM HEPES, pH 7.4, 200 mM NaCl, 2 mM MgCl_2_, 1% (vol/vol) Triton X-100, 0.5 mM TCEP, protease inhibitors (AEBSF, Leupeptin and Benzamidine) and supplemented with phosphatase inhibitors (50 mM NaF and 10 mM beta-glycerophosphate) at 4 °C. Lysates were clarified by centrifugation (15,000 g for 40 min at 4 °C) and incubated with 5 mL glutathione Sepharose 4B (GE Healthcare) for 1.5 hr at 4 °C with gentle shaking. The glutathione Sepharose 4B matrix was applied to a gravity column, washed with 100 mL wash buffer (20 mM HEPES, pH 7.4, 200 mM NaCl, 2 mM MgCl_2_, and 0.5 mM TCEP), and purified complexes were eluted with 40 mL wash buffer containing 50 mM reduced glutathione. Eluted complexes were treated with TEV protease at 4 °C overnight. TEV-treated complexes were purified to homogeneity by injection on Superose 6 Increased 10/300 GL (GE Healthcare) column that was pre-equilibrated in gel filtration buffer (20 mM HEPES, pH 7.4, 200 mM NaCl, 2 mM MgCl_2_, and 0.5 mM TCEP). For long-term storage, fractions from the gel filtration chromatography were snap frozen in liquid nitrogen and kept at -80 °C.

For expression of His_6_-tagged ARF1 (residues E17-K181) and all the ARF1 mutants, the plasmids were transformed into *E. coli* BL21 DE3 star cells and induced with 1 mM IPTG for 3 hr. The cells were lysed in 50 mM Tris-HCl pH 8.0, 300 mM NaCl, 2 mM MgCl_2_, 10 mM imidazole, 0.5 mM TCEP and 1 mM PMSF by ultrasonication. The lysate was centrifuged at 15,000 x g for 30 min. The supernatant was loaded into Ni-NTA resin and washed with 20 and 50 mM imidazole and eluted with 300 mM imidazole. The eluate was further purified on a Superdex 75 10/300 GL (GE Healthcare) column equilibrated in 20 mM HEPES, pH 7.4, 200 mM NaCl, 2 mM MgCl_*2*_, and 0.5 mM TCEP. STREP-STREP-FLAG tag RAB8A (residues M1-S181) was expressed in HEK293-GnTi cells and purified by Strep resin and eluted in 10 mM desthiobiotin buffer. The eluted protein was applied on Superdex 75 10/300 GL column. The nucleotide binding state of each small GTP-binding protein was assessed by HPLC (see below). The purified RAB8A was GDP bound and our previous data demonstrated that it had spontaneous guanine exchange activity ^21^, therefore RAB8A was incubated with GTP at room temperature for 2 hrs and applied to a PD 10 column to remove the excess nucleotide. The resulting RAB8A sample had roughly 45% GTP bound. ARF1^I49H/G50R^ and ARF1 ^I49H/G50R/K73W^ were also GDP bound, therefore the GTP reloading protocol was also applied to these samples.

For the assembly of C9orf72:SMCR8:WDR41 with ARF1^Q71L^ complex, 15 μM of C9orf72:SMCR8:WDR41 was incubated with 15 fold of ARF1^Q71L^ in 500 μl overnight and injected into Superose 6 Increased 10/300 column.

### Cryo-electron microscopy grid preparation and data imaging

For the Talos Arctica dataset, the purified complex was diluted to 0.16-0.2 mg/ml in 20 mM HEPES pH 7.4, 200 mM NaCl, 2 mM MgCl_2_ and 0.5 mM TCEP, and incubated with BeF_3^-^_overnight. 3.5 μL of sample was deposited onto freshly glow-discharged (PELCO easiGlow, 30 sec in air at 20 mA and 0.4 mbar) C-flat 1.2/1.3-3Au grids (Electron Microscopy Sciences), and blotted for 4 sec using a Vitrobot Mark IV (FEI) with 30 sec incubation, blot force 8 and 100 % humidity and temperature set to 22°C. The complex was visualized in the Talos Arctica operated at an acceleration voltage of 200 kV, and equipped with a Gatan K3 Summit director electron detector in super-resolution counting mode at 36,000x, corresponding to a pixel size of 0.57 Å. In total, 2,579 movies were acquired with the defocus range between -1 to -2.5 μm with a 3 x 3 image shift pattern prior to stage movement. All movies consisted of 50 frames with a total dose of 50 e^-^/Å^2^ and a total exposure time of 5.16 sec.

For the untilted Titan Krios dataset, grids were prepared in the same manner and the complex was collected using a Titan Krios G2 electron microscope (FEI) operating at 300 kV with a Gatan Quantum energy filter (operated at 20 eV slit width) using a K3 summit direct electron detector (Gatan, Inc.) in super-resolution counting mode at magnification 81,000x, corresponding to a pixel size of 0.47 Å on the specimen level. In total, 10,188 movies were collected with defocus of -1.2 to -2.5 μm with a 3 x 3 image shift pattern prior to stage movement. Movies consisted of 50 frames, with a total dose of 50 e-/Å^2^ and a total exposure time of 2.929 sec.

The Titan Krios dataset with a stage tilt of 30° was obtained using 4 μL sample applied on freshly glow-discharged (PELCO easiGlow, 25 sec in air at 25 mA and 0.4 mbar) UltrAufoil R1.2/1.3 grid (Ted Pella) and plunged into liquid ethane after 30 sec incubation and blotting for 4 sec using a Vitrobot Mark IV (FEI) with blot force 0 and 100 % humidity and temperature set to 5°C. 4,585 movies were collected using SerialEM at around -1.5 to 2.8 μm defocus with 1 x 3 image shift pattern. The movies were exposed to a total dose of 50 e-/Å^2^ for 5.326 sec at CDS mode.

The Titan Krios dataset with a stage tilt of 40° was collected using 4 μL a sample applied on freshly glow-discharged (PELCO easiGlow, 25 sec in air at 25 mA and 0.4 mbar) GF-1.2/1.3-3Au-45nm-50 grid (Electron Microscopy Sciences) and plunged into liquid ethane after 30 sec incubation and blotting for 3 sec using a Vitrobot Mark IV (FEI) with blot force 8 and 100 % humidity and temperature set to 4°C. 6,178 movies were collected using SerialEM at -1.5 to 2.8 μm defocus with 1 x 3 image shift pattern. The movies were exposed to a total dose of 50 e-/Å^2^ for 5.326 sec at CDS mode.

All of the data were acquired with SerialEM using custom macros for automated single particle data acquisition. Imaging parameters for the data set are summarized in Extended Data Table 1.

### Cryo-electron microscopy data processing

For the Talos Arctica dataset, drift and beam-induced motion correction was applied on the super-resolution movie stacks using MotionCor2 ^44^ and using 7 x 5 patches and binned 2x by Fourier cropping to a calibrated pixel size of 1.14 Å/pixel. The defocus values were estimated by CTFFIND4 ^45^ from summed imaged without dose weighting. After manual check to remove empty or crystalline ice micrographs, 251 micrographs were discarded and the remaining 2,328 micrographs were subjected to reference-free particle picking using gautomatch-v0.53 (http://www.mrc-lmb.cam.ac.uk/kzhang/Gautomatch/). About 4 million particles were extracted with binning 4x in Relion 3.1 ^46^ and applied to iterations of three-dimensional (3D) classifications. The best class was selected and re-extracted with binning 1x. The particles at the edges were removed and imported into cryoSPARC v2 ^47^ for further two-dimensional (2D) classification clean up and non-uniform refinement. The resulting 80,569 particles were refined to 4.84 Å and used as reference map for further Titan Krios dataset processing. The work-flow is depicted in Extended Data Fig 2.

For the untilted Krios dataset, drift and beam-induced motion correction was applied on the super-resolution movie stacks using MotionCor2 using 7 x 5 patches and binned 2x to a calibrated pixel size of 0.94 Å/pixel. The defocus values were estimated by CTFFIND4 from summed imaged without dose weighting. Other processing procedures were performed within Relion 3.1. 7,347 micrographs with CtfMaxResolution values better than 5 were selected for further processing. Around 6 millions of particles were picked using gautomatch-v0.53 (http://www.mrc-lmb.cam.ac.uk/kzhang/Gautomatch/) and then extracted with binning 6x. The particles were subjected to 3D classification using a 45 Å low-pass filtered reference generated by the reconstruction collected in Talos Arctica microscope. 888,231 particles from the best class was selected and refined to 7.755 Å. The refined particles were further cleaned up with rounds of 2D classification and 3D classification. The extracted, unbinned particles was refined to 4.5 Å and applied to CTF refinement for beam-tilt and per-particle defocus determination as well as Bayesian polishing. The shiny particles was further refined to 3.7 Å. Map sharpening was performed with DeepEMhancer ^48^ using the “tightTarget” or “highRes” models. The work-flow is depicted in Extended Data Fig.3.

For the Krios dataset with a stage tilt of 30°. data processing followed a similar pipeline of 0° tilt dataset for processing except that first run a full-micrograph based CTF estimation, then pick particles using gautomatch and then subjected the particle coordinates to re-run local CTF estimation on a per-particle based in gctf ^49^ (Extended data Fig. 3). For the Krios dataset with a stage tilt of 40°, the data was imported into cryoSPARC v2 for patch motion correction and patch CTF estimation. Particle picking was done using 2D class averages from 0° tilt as reference for Autopick in Relion3.1 and subjected to several rounds of 3D classification. The best classes were imported into cryoSPARC v2 for further rounds of 2D classification. The clean particles were applied to both global and local CTF refinement for then imported into Relion 3.1 to merge with other two datasets (Extended data Fig. 3).

All reported resolutions are based on the gold-standard FSC 0.143 criterion. cryoSPARC v2 database files were converted to Relion star file using the UCSF PyEM package script (https://github.com/asarnow/pyem/). Particle orientation distributions were plotted using scripts (https://github.com/fuzikt/starpy). 3DFSC curve for the reconstruction was calculated on the 3DFSC processing server ^50^.

### Atomic model building and refinement

For the C9orf72:ARF1-SMCR8: WDR41 model, one copy of the coordinates from all three subunits (PDB 6V4U or 6LT0) ^20,21^, WDR41 (PDB 6WHH) and mouse ARF1 (PDB: 1O3Y) ^31^ were rigid body fitted separately into the density map using UCSF Chimera ^51^. The Leu71 of ARF1 was mutated to its wild type residue Gln and the GTP was replaced as GDP-BeF_3^-^_. Atomic coordinates were refined by iteratively performing Phenix real-space refinement and manual building and correction of the refined coordinates in Coot. To avoid overfitting, the map weight was set to 1 and secondary structure restraints were applied during automated real-space refinement. Model quality was assessed using MolProbity ^52^ and the map-vs-model FSC (Extended Data Table 1, Extended Data Fig. 6b). A half-map cross-validation test showed no indication of overfitting (Extended Data Fig.6a). Figures were prepared using UCSF Chimera. The cryo-EM density map has been deposited in the Electron Microscopy Data Bank under accession code EMD-23827 and the coordinates have been deposited in the Protein Data Bank under accession number 7MGE.

### High-performance liquid chromatography analysis of nucleotides identity bound to small GTPases

The nucleotides bound to small GTPases were assessed by denaturing 30 μl of protein to 95 °C for 5 min followed by 5 min centrifugation at 16,000 x g. The supernatant was loaded onto a HPLC column (Eclipse XDB-C18, Agilent). Nucleotides were eluted with HPLC buffer (10 mM tetra-n-butylammonium bromide, 100 mM potassium phosphate pH 6.5, 7.5 % acetonitrile) at flow rate 0.6 ml/min. The identity of the nucleotides was compared to GDP and GTP standards.

### HPLC-based GAP assay

HPLC-based GTPase assays were carried out by incubating 30 μl of 30 μM GTPases with or without GAP complex at a 1:100, 1:50 or 1:10 molar ratio as indicated for 15 min at 37 °C. Samples were immediately boiled for at least 5 min at 95 °C and centrifuged for 5 min at 16,000 g. The supernatant was injected onto an HPLC column as described above. The experiments are carried out at least three times and one representative plot is shown.

---

**Extended Data Fig 1:**
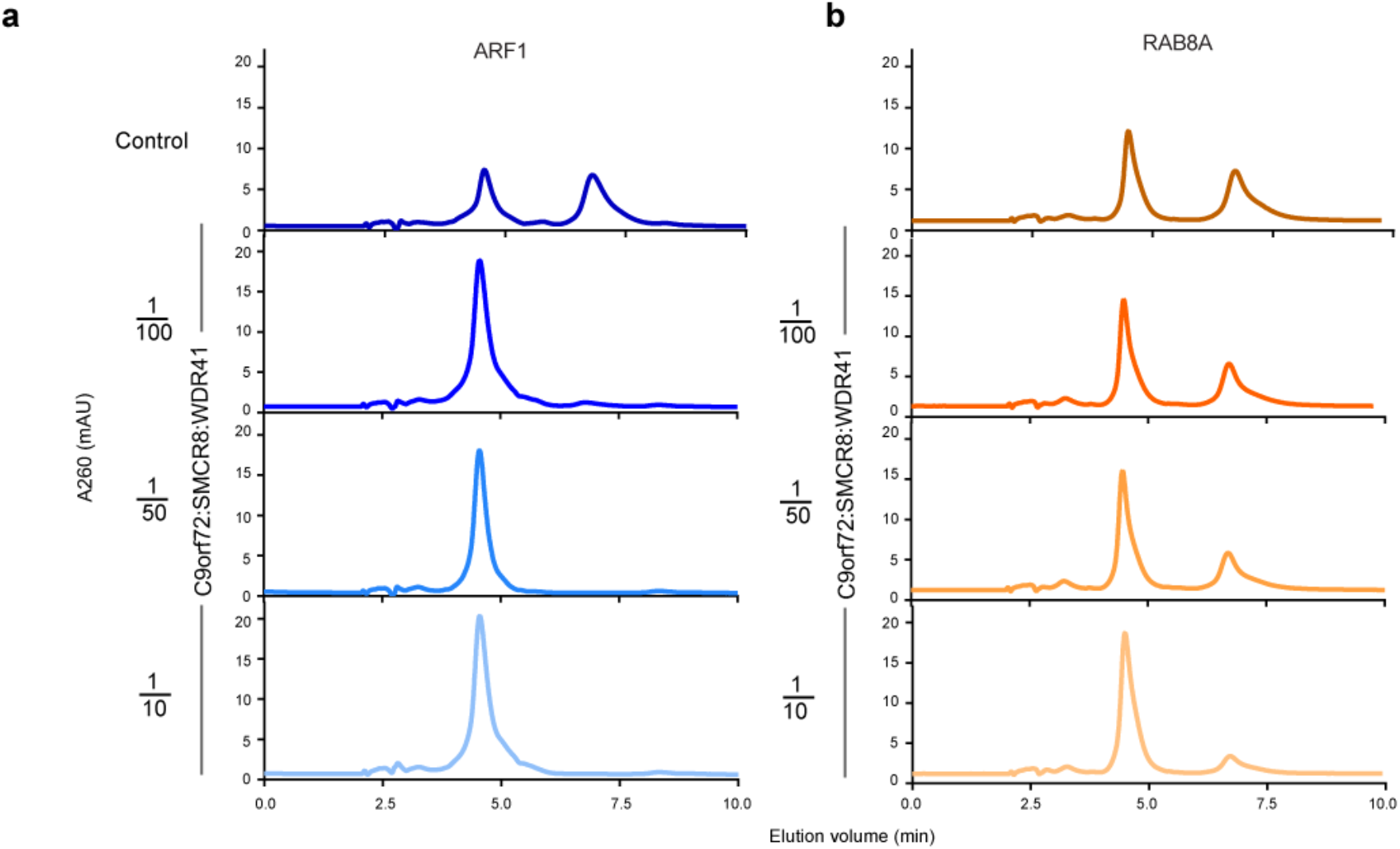
The HPLC result of incubating ARF1 and RAB8A with different molar ratio of C9orf72:SMCR8:WDR41 complex. 30 µM of ARF1 (a) or RAB8A (b) was treated with 0, 0.3, 1.5 and 3 µM C9orf72:SMCR8:WDR41 complex for 15 min at 37°C.

**Extended Data Fig 2:**
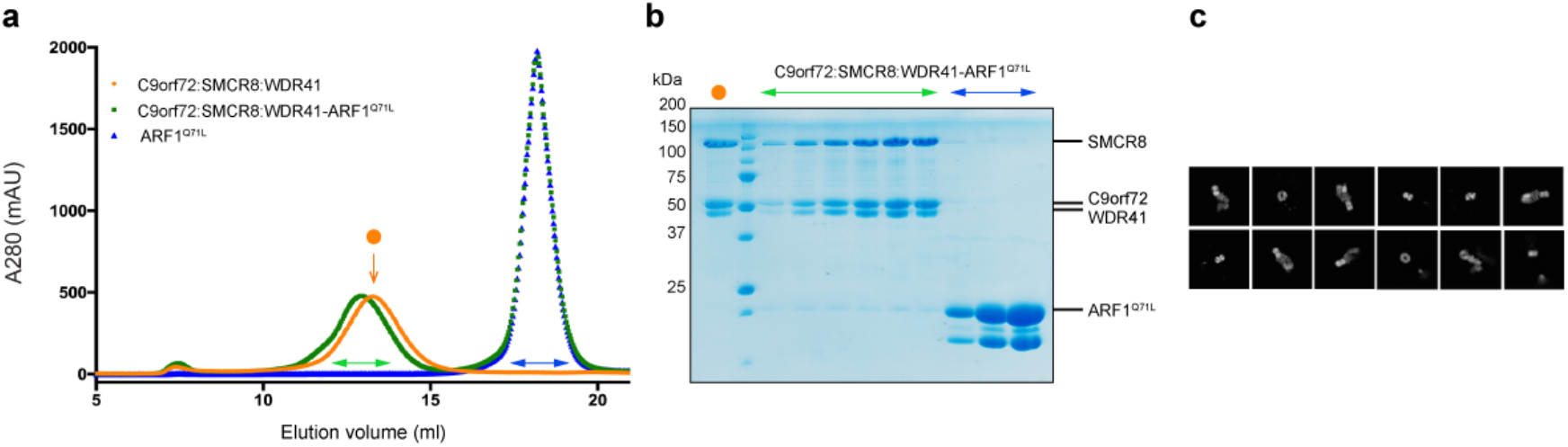
Assembly of C9orf72:SMCR8:WDR41:ARF1^Q71L^. a, the Superose 6 size exclusion profile of the reconstituted C9orf72:SMCR8:WDR41:ARF1^Q71L^ complex. b, the SDS-PAGE analysis of the peak fractions. c. 2D class averages for the C9orf72:SMCR8:WDR41:ARF1^Q71L^ complex.

**Extended Data Fig 3.**
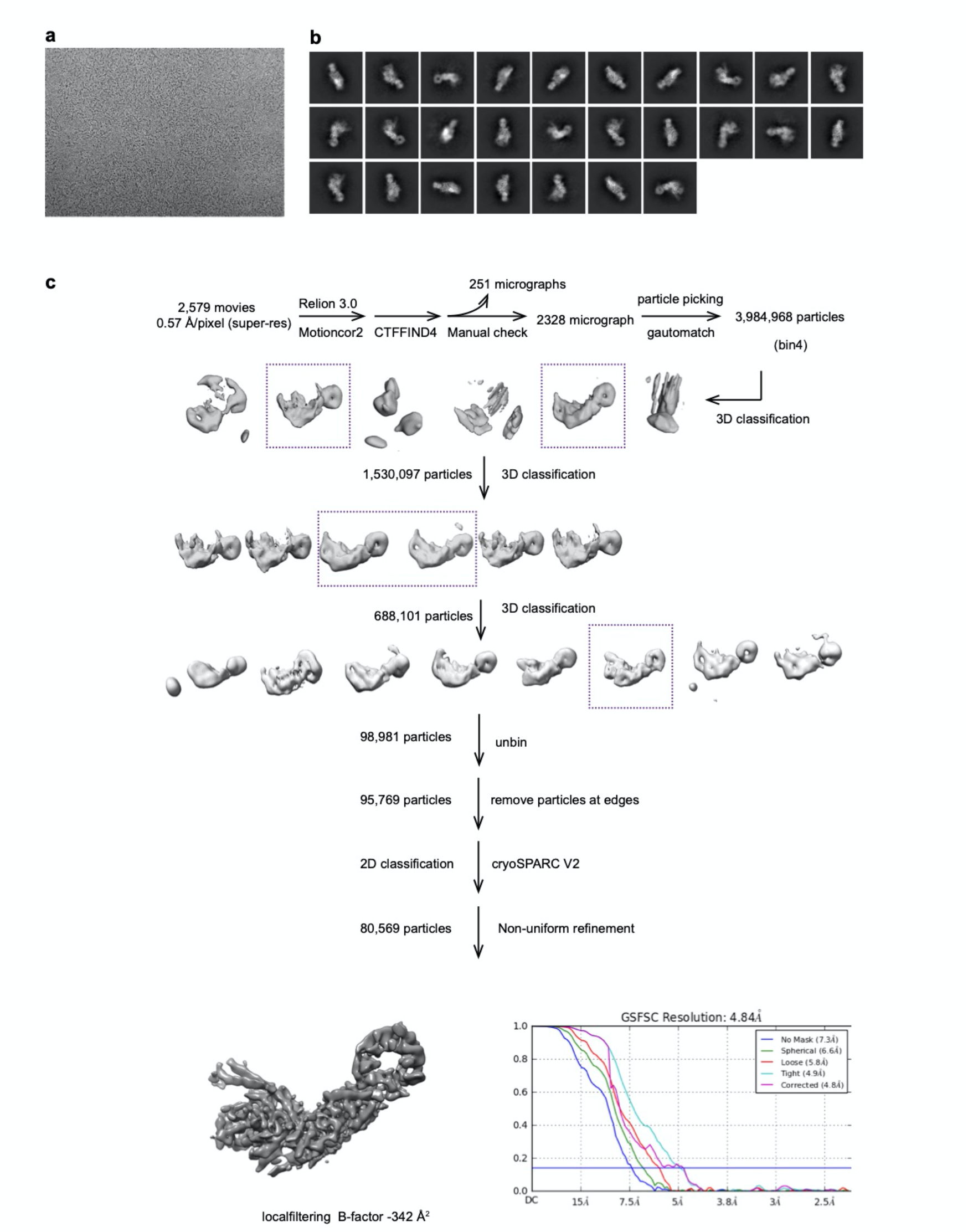
Cryo-EM data processing work flow for C9orf72:ARF1-SMCR8:WDR41 complex incubated with BeF_3_^-^in 200 kV Arctica microscope. a, Representative cryo-EM micrograph of C9orf72:ARF1-SMCR8:WDR41 complex. b, Representative 2D classes. c, Image processing procedure for the dataset collected at 200 kV Talos Arctica.

**Extended Data Fig 4:**
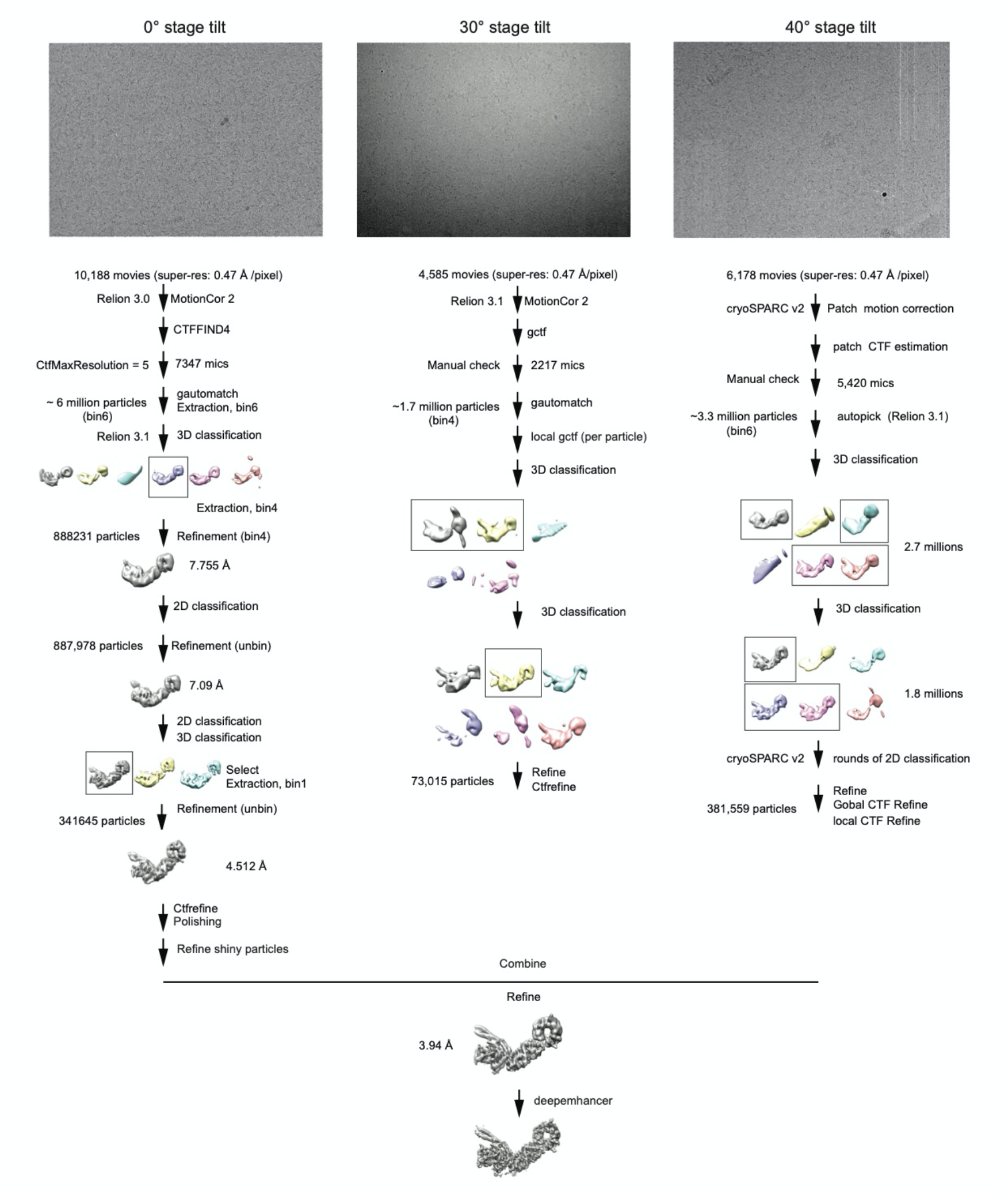
**Cryo-EM data processing work flow for C9orf72:ARF1-SMCR8:WDR41 complex incubated with BeF_3_^-^ in 300 kV Titan Krios.**

**Extended Data Fig 5:**
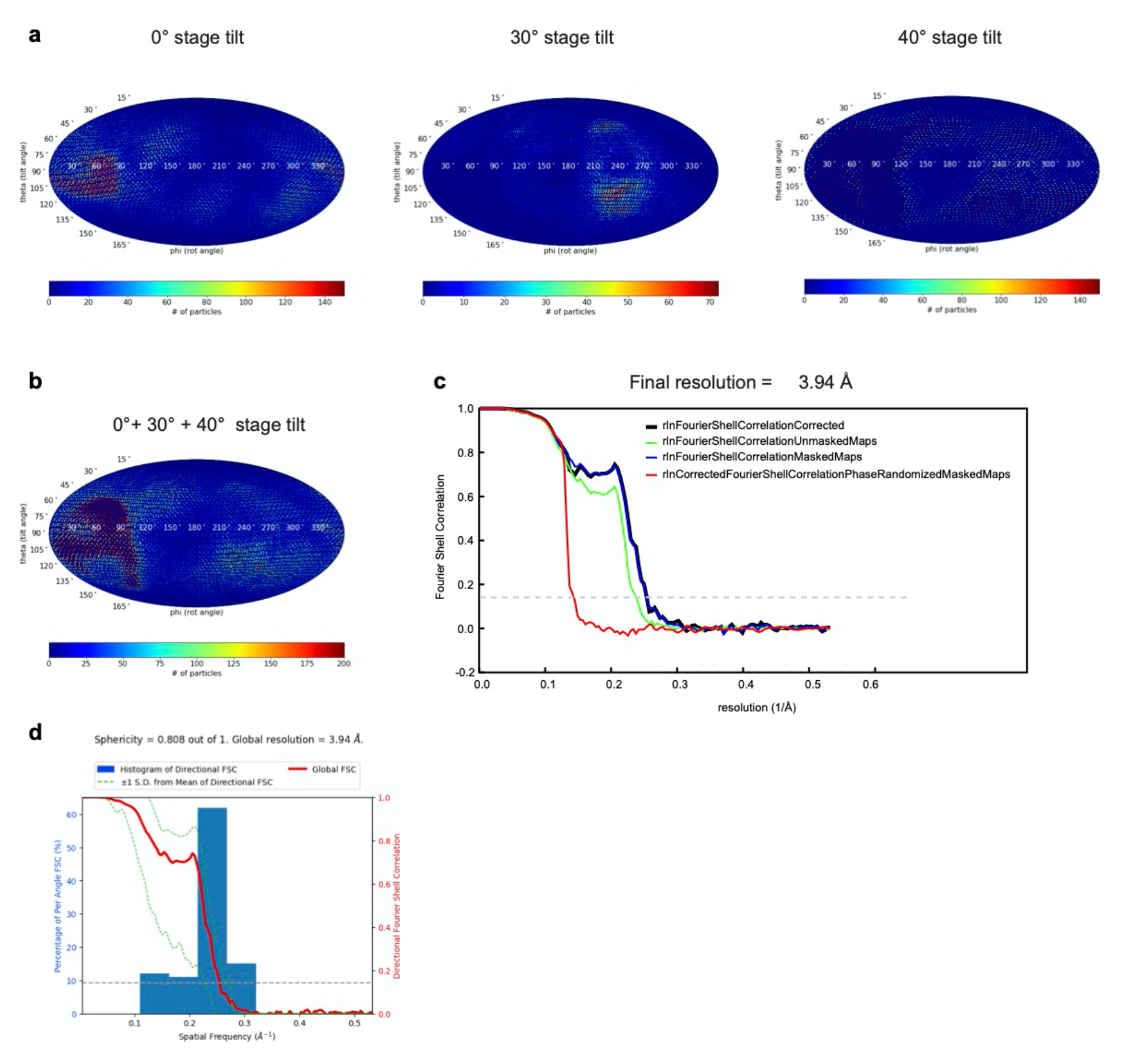
Angular distribution of the particles in the reconstructions. a-b, heatmap of the particle orientations from the 0°, 30° and 40° tilted datasets (a) or combined datasets (b) shown in Mollweide representations. c, Comparison between the FSC curves. d, 3DFSC plot for the final reconstruction.

**Extended Data Fig 6:**
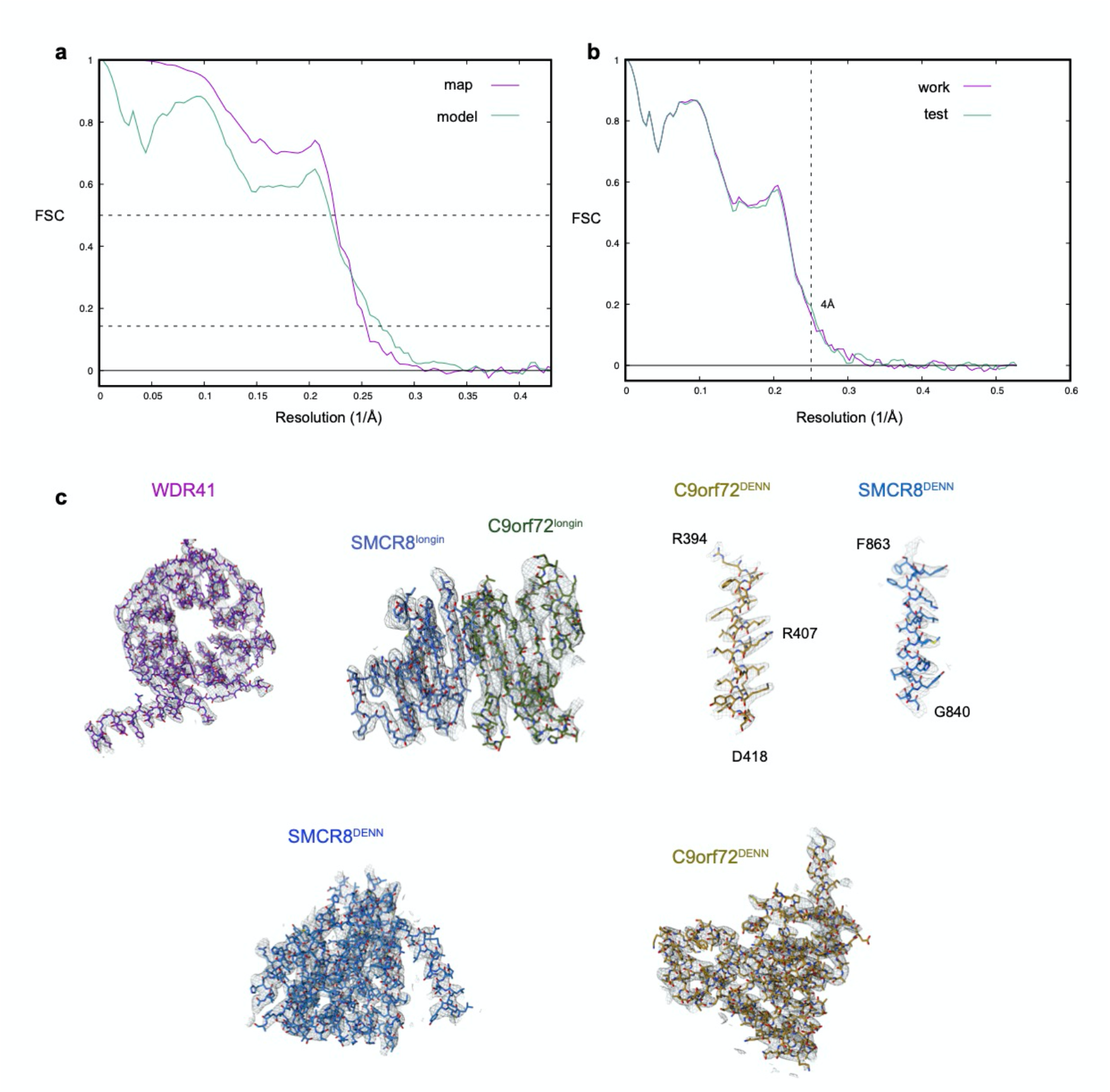
Model building and validation. a, Refinement and map-vs-model FSC. B, Cross-validation test FSC curves to assess over-fitting. The refinement target resolution of 4 Å is indicated c, Refined coordinate model fit of the indicated region in the cryo-EM density.

**Supplementary Fig 1:**
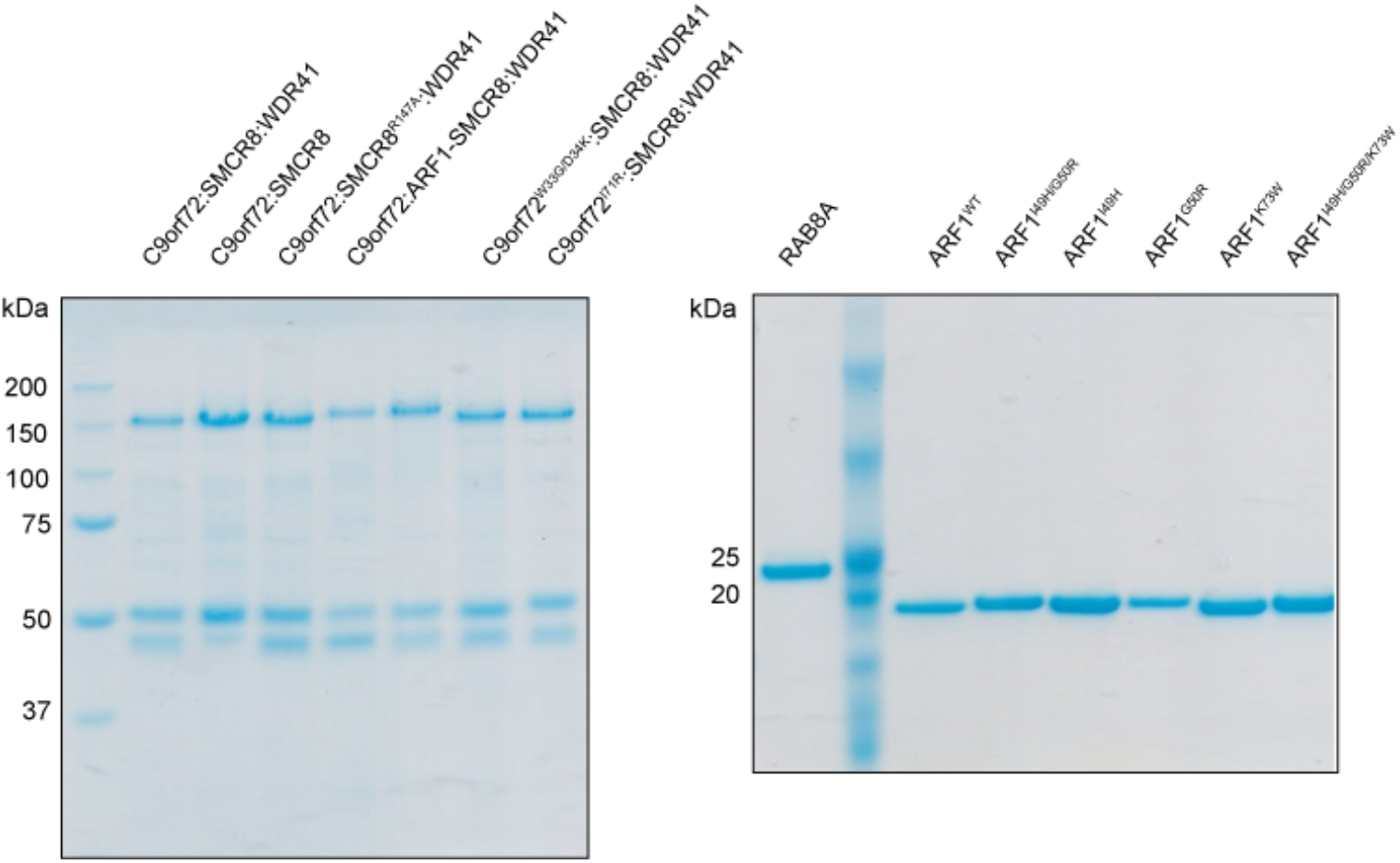
SDS-PAGE analysis of the purified proteins used is this study.

**Extended Data Table 1:** Cryo-EM data collection, refinement and validation statistics.

|  |  |
| --- | --- |
|  | C9orf72:ARF1-SMCR8:WDR41 (EMD-23827)<br>(PDB 7MGE) |
| **Data collection and processing** |  |
| Microscope | Titan Krios |
| Magnification (calibrated) | 43,516 |
| Camera | Quantum-K3 Summit |
| Voltage (kV) | 300 |
| Electron exposure (e<sup>-</sup>/Å<sup>2</sup>) | 50 |
| Pixel size (Å) | 0.94 |
| Symmetry imposed | C1 |
| Initial particle images (no.) | ~11 million |
| Final particle images (no.) | 796,219 |
| Map resolution (Å) | 3.94 |
| FSC threshold | 0.143 |
| **Refinement** |  |
| Initial model used (PDB code) | 1O3Y, 6V4U, 6LT0, 6WHH |
| Map sharpening *B* factor (Å<sup>2</sup>) | - |
| Model composition |  |
| Non-hydrogen atoms | 9017 |
| Protein residues | 1323 |
| Ligands | GDP, BeF<sub>3</sub><sup>-</sup>, Mg<sup>2+</sup> |
| R.m.s. deviations |  |
| Bond lengths (Å) | 0.003 |
| Bond angles (°) | 0.687 |
| Validation |  |
| MolProbity score | 2.16 |
| Clashscore | 10.98 |
| Poor rotamers (%) | 0 |
| Ramachandran plot |  |
| Favored (%) | 88.04 |
| Allowed (%) | 11.48 |
| Disallowed (%) | 0.48 |

